# Colistin resistance-mediated lipopolysaccharide modification in *Klebsiella pneumoniae* modulates host inflammatory response

**DOI:** 10.64898/2026.07.30.741922

**Authors:** Avijit Dutta, Peter Gallagher, Katrien M.J. Sutherland, Borna Halder, To Nguyen Thi Nguyen, Jacqueline A. Keane, Gerald Larrouy-Maumus, Stephen Baker

**Affiliations:** Cambridge Institute of Therapeutic Immunology and Infectious Disease, Department of Medicine, University of Cambridge, Cambridge, United Kingdom; Department of Microbiology and Veterinary Public Health, Chattogram Veterinary and Animal Sciences University, Chattogram-4225, Bangladesh; Department of Statistics, University of Chittagong, Chittagong, Bangladesh; Department of Microbiology, Faculty of Medicine, Nursing and Health Sciences, Monash University, Melbourne, VIC 3800, Australia; MRC Centre for Molecular Bacteriology and Infection, Department of Life Sciences, Faculty of Natural Sciences, Imperial College London, London, United Kingdom

**Author notes:** Corresponding author, (AD). School of Chemistry, University of Edinburgh, Edinburgh, United Kingdom (Katrien M.J. Sutherland). A*STAR Infectious Diseases Labs (A*STAR IDL), Agency for Science, Technology and Research (A*STAR), Singapore 138648, Singapore.

**Keywords:** colistin resistance, lipopolysaccharide, *K. pneumoniae*, immunity

## Abstract

Resistance to the polymyxin antimicrobial colistin in Gram-negative bacteria is associated with a modification of the immunogenic lipid A moiety of the lipopolysaccharide (LPS). Chromosomal and plasmid-borne colistin resistance results in the addition of L-Ara4N and pEtN groups to lipopolysaccharide (LPS), respectively. Here, using THP-1 cells, we studied the impact of different LPS modifications of *Klebsiella pneumoniae* in stimulating host immune response. *K. pneumoniae* clinical isolates were screened for colistin resistance using broth microdilution (BMD) and the MALDIxin test. LPS was extracted from colistin-resistant isolates and used to stimulate differentiated THP-1 cells. Luminex cytokine assay measured the immune induction via a panel of proinflammatory cytokines. Out of a collection of 72 clinical *K. pneumoniae*, eight (11.1%) exhibited phenotypic colistin resistance with a minimum inhibitory concentration (MIC) of 8 to 64 mg/L. In total, five isolates possessed genes associated with polymyxin resistance; three isolates had a mutation in the *pmrB* gene, and two were *mcr-8.1* positive. MALDIxin demonstrated that all eight phenotypic colistin-resistant isolates elaborated peaks at m/z 1,955 and m/z 2,193, indicating an L-Ara4N group of LPS modification. For two *mcr-8.1* positive isolates, LPS had a pEtN group. The LPS modification positively correlated with colistin MIC (correlation coefficient, r= 0.6 and R^2^= 0.4). Compared to the native structure, LPS modification was associated with greater production of IL-1β, IL-6, and CXCL-8 (*p*<0.001). The pEtN-conjugated LPS triggered a significantly greater production of TNF-α, IL-6, and CXCL-8 compared to L-Ara4N (*p*<0.05). This study reveals that the colistin MIC value can significantly predict lipid A modification in clinical *K. pneumoniae,* and differences in resistance-mediated lipid A modification result in variation in the immunological response. This study highlights the potential of dynamic host-pathogen interaction in the context of colistin resistance.

## Introduction

In the era of antimicrobial resistance (AMR), polymyxin E (colistin) is regarded as one of the key last-line antimicrobials against pan-drug-resistant bacterial infections [1]. KPC-type carbapenemase-producing *K. pneumoniae* is an emerging problem in healthcare settings and often requires the administration of colistin [2]. In addition to having a bactericidal action, colistin also has an endotoxin-neutralisation function [3]. In recent years, an upward trend of colistin resistance has been reported in *K. pneumoniae,* with a resistance prevalence of 4.8% in 2013 and increasing to 8.2% in 2022 [4,5].

Colistin resistance in *K. pneumoniae* has been characterised primarily by chromosomal mutations leading to the addition of L-Ara-4N to LPS [6]. Alternatively, plasmid-borne colistin resistance, which has increased significantly in recent years, is encoded by *mcr* genes, leading to the development of PEtN-conjugated LPS [7,8]. Some studies have also reported other colistin resistance mechanisms, such as palmitoylation and/or hydroxylation of lipid A, in *K. pneumoniae* [9,10]. More recently, mass spectrum data have been successfully used not only to detect colistin resistance but also to characterise the extent of lipid A modification and to distinguish chromosomal and plasmid-borne resistance mechanisms [6,11].

In Gram-negative bacteria, LPS is a pathogen-associated molecular pattern (PAMP) that activates innate immunity pathways. Modification of lipid A shifts the host immune response during infections [12]. Studies have established that expression of the plasmid-borne colistin resistance gene, *mcr*, is associated with increased pathogen virulence and higher mortality [13]. Some studies suggest that lipid A modification due to colistin resistance results in the greater production of pro-inflammatory cytokines and immune checkpoints [14,15]. Compared to the wild-type, colistin-resistant *K. pneumoniae* show enhanced intracellular killing evasion, higher monocyte apoptosis, and increased production of TNF-α, IL-6 and CXCL10 [14]. However, contentious findings exist, and it has been reported that LPS of *mcr*-positive isolates induces lower IL-6 and TNF-α compared to control LPS [15].

There is limited data regarding LPS modification derived from colistin resistance and how it may impinge on the immune response during infection. The impact of different types and degrees of lipid A modification on colistin’s anti-endotoxin ability has not been explored.

Here, we exploited mass spectrum data to determine lipid A modification of the colistin-resistant *K. pneumoniae* isolates and measured its relationship with colistin MIC. Subsequently, using an *in vitro* THP-1 cell line, we investigated the immunological impact of different degrees and variants of LPS modification attributed to colistin resistance.

## Materials and methods

### K. pneumoniae isolates

The *K. pneumoniae* isolates used in this study were obtained from a previously conducted retrospective study at the Hospital for Tropical Diseases and Oxford University Clinical Research Unit, Ho Chi Minh City, Vietnam. Bacteria were isolated from clinical samples from patients admitted at the Hospital for Tropical Diseases, Ho Chi Minh City, Vietnam, between 2010 and 2013. Patients had different clinical conditions, predominantly septicaemic, hepatic disorders and HIV infection. A clinical sample was collected from each case, and then blood culture was performed following standard bacteriological procedures. The patient data were accessed during the previous study period of April 2014 to December 2015 for research purposes without the possibility of identifying individual patients. The isolates were stored at -80°C, genome sequenced, and later transferred to the Cambridge Institute of Therapeutic Immunology and Infectious Diseases, Cambridge, United Kingdom.

### MALDI-TOF and BMD assays

*K. pneumoniae* species was confirmed using matrix-assisted laser desorption/ionization-time of flight (MALDI-TOF) spectrometry following the established protocol described previously [18]. The MIC of the *K. pneumoniae* isolates against colistin was determined using the BMD method. In the BMD assay, *E. coli* ATCC 25922 was used as a positive control. Each isolate was studied in triplicate, and the results were recorded and interpreted following CLSI standards [19]. A colistin MIC of ≥4 mg/L was considered resistant.

### Detection of colistin resistance by MALDIxin test

*K. pneumoniae* isolates that were phenotypically resistant to colistin were subjected to MALDIxin (S1 Fig) [11]. Samples were analyzed at 20 kV using the negative ion mode of this MALDI-TOF system with an extraction delay time of 20 ns. Each sample was conducted in triplicate. Mass spectrometric analysis-based colistin resistance was studied using the reflectron mode of the 4,800 Proteomics Analyzer (Applied Biosystem, USA). At first, the intensities of the peaks for naïve LPS (m/z 1,824 and m/z 2,062) and modified LPS (m/z 1,955 and m/z 2,193) were measured [11]. To calculate the %LPS modification, the total intensity of modified LPS was divided by the sum of modified and unmodified LPS intensities and expressed as a percentage. To determine the type of modification, a shift of m/z +123 and m/z +131 to the naïve LPS intensities was indicated as the pEtN and L-Ara4N modifications of lipid A, respectively [11].

### Extraction and quantification of LPS

*K. pneumoniae* isolates of different LPS modifications were selected for the cell culture experiment. To extract LPS from colistin-resistant *K. pneumoniae*, isolates were cultured on LB agar supplemented with 4 mg/L colistin. The LPS was extracted using the protocol described by a commercial LPS isolation kit (MAK339, Sigma-Aldrich, USA). Extracted LPS was quantified using Pierce^TM^ LAL Chromogenic Endotoxin Quantitation Kit (Thermo Scientific, USA). Here, the blank-corrected absorbance of known *E. coli* endotoxin standards was used to generate a standard curve. Finally, the sample LPS quantity was determined from the observed absorbance (A_405nm_) using a linear regression plot.

### In-vitro THP-1 cell culture experiment

Human monocyte-derived THP-1 cells (ATCC TIB-202^TM^) were used to study the effect of colistin resistance-mediated *K. pneumoniae* lipid A modifications on the host immune system following the protocol described earlier [14]. The detailed workflow is outlined in the supplementary information (S2 Fig). Briefly, cell culture media were prepared with RPMI 1,640 medium (450 ml), foetal bovine serum (50 ml), sodium pyruvate (5 ml), and N-2-hydroxyethylpiperazine-N’-2-ethanesulfonic acid (5 ml). Regular passaging and counting of the cells were performed. Three days prior to LPS exposure, the THP-1 cells were differentiated using phorbol 12-myristate 13-acetate (PMA) (1:10,000). The morphology of THP-1 cells was observed under a microscope before and after each splitting and differentiation. The differentiated THP-1 cells were stimulated with 10 µl of LPS at a concentration of 10 ng/ml. LPS from *E. coli* O111:B4 (Sigma-Aldrich, USA) and endotoxin-free water were used as positive and negative controls, respectively. After overnight incubation, supernatant was collected, labelled and stored at -80°C for the Luminex cytokine assay. In addition, how LPS modification affects the endotoxin neutralisation ability of colistin was studied following the protocol described by Roberts *et al*., with some modifications [20]. All the samples were tested in duplicate.

### Luminex cytokine assay

A panel of pro-inflammatory cytokines (TNF-α, IL-1β, IL-6, IL-10 and CXCL-8) were investigated in the differentiated THP-1 cell culture supernatant using Bio-Plex Pro Human cytokine assay kit following the manufacturer’s instructions. The plate reading and data acquisition were performed on the Bio-Plex 2200 system (Bio-Rad, USA). The system generated separate standard curve plots for each assay and measured the cytokines in each sample. The data were plotted using GraphPad Prism (Boston, USA).

### Kleborate analysis

Genomic characterisation of the *K. pneumoniae* isolates was obtained using the Kleborate V2.1.0 pipeline [21]. The analysis resulted in species assignment, sequence typing, and detection of resistance and virulence genes. The pipeline profiled the isolates based on virulence and resistance score on the availability of clinically important gene markers. The resistance scoring system was based on the genes associated with ESBL only (score 01), carbapenemase only (score 02), and carbapenemase plus colistin resistance (score 03). Virulence scores range from 0 to 5; no virulence gene (score 0), yersiniabactin only (score 01), colibactin without aerobactin (score 02), aerobactin only (score 03), aerobactin and yersiniabactin without colibactin (score 04), and all three genes (score 05).

### Pangenome and phylogenetic analysis

The *K. pneumoniae* sequence data were assembled using the Global Health Research Unit (GHRU) assembly pipeline v2.1.2 and annotated with Prokka v1.14.6 [22]. The pangenome analysis of the. *K. pneumoniae* study isolates was conducted using the Roary pipeline v3.12 [23]. The minimum BLASTP identity was set at 90%, and a 1.5 inflation value for the Markov clustering technique. The SNP-based phylogenetic analysis of the WGS was performed using pipelines from GHRU on Nextflow [24,25]. After quality checking by MultiQC, the reads were mapped to the reference of the *K. pneumoniae* genome (accession CP064352.1) using bactmap v0.9. Default parameters were selected for the analysis unless otherwise specified. A maximum likelihood phylogenetic tree was generated from single-nucleotide polymorphism (SNPs) alignment using IQTREE v1.6.10 and visualised with iTOL, v5 [26,27].

The tree was plotted with sequence type (ST), K and O loci type, resistance and virulence score, AMR gene number, ESBL gene and plasmid type. Other information, such as AMR phenotype, mucoviscosity, and year of isolation, was added to the phylogenetic tree.

## Results

### Characteristics of the clinical cases

The *K. pneumoniae* isolates were genome sequenced and grouped into virulent *K. pneumoniae* (Vir-Kp) and non-virulent *K. pneumoniae* (non-Vir-Kp) via the presence/absence of yersiniabactin (*Ybt*), colibactin (*Clb*), and aerobactin (*Abt*) (Table 1). Hepatic illness, septicaemia and HIV infections comprised >60% of the total *K. pneumoniae* infections. Out of 19 hepatic cases, 13 patients suffered from cirrhosis, and 10 were associated with Vir-Kp (12.1%). More than 16% of the patients with either septicaemia or AIDS had an infection with Vir-Kp. Out of all cases, 11 were associated with hypervirulent *K. pneumoniae* isolates (virulence score 5). Among the hypervirulent *K. pneumoniae*, a higher prevalence of antimicrobial resistance was observed against sulfamethoxazole-trimethoprim (19.3%), ceftriaxone (18.1%), ceftazidime (18.1%) and ticarcillin-clavulanic acid (16.9%). The isolates were generally susceptible to amikacin, imipenem, meropenem, and ertapenem with 2.4%, 2.4%, 3.6%, and 7.2% resistance, respectively.

**Table 1:** Characteristics of clinical cases and AMR pattern of *K. pneumoniae* study isolates.

| Case status, isolation year and antimicrobial resistance profiles | Case no (%) |  |
| --- | --- | --- |
|  | Vir-Kp<sup>a</sup> | Non-Vir-Kp<sup>b</sup> |
| **Clinical cases** |  |  |
| Urinary tract infections (UTI) | 1 (1.2%) | 3 (3.6%) |
| Septicaemia | 7 (8.4%) | 11 (13.3%) |
| Viral infections | 5 (6.0%) | 3 (3.6%) |
| Hepatic diseases (cirrhosis, hepatitis) | 10 (12.1%) | 9 (10.9%) |
| AIDS/HIV infection | 7 (8.4%) | 8 (9.6%) |
| Pneumonia and respiratory failure | 2 (2.4%) | 1 (1.2%) |
| Tetanus | 3 (3.6%) | 3 (3.6%) |
| Others | 3 (3.6%) | 7 (8.4%) |
| **Isolation year** |  |  |
| 2010 | 9 (10.9%) | 4 (4.8%) |
| 2011 | 10 (12.1%) | 8 (9.6%) |
| 2012 | 9 (10.9%) | 9 (10.9%) |
| 2013 | 6 (7.2%) | 10 (12.1%) |
| 2014 | 9 (10.9%) | 9 (10.9%) |
| **Antimicrobial resistance** |  |  |
| Amoxicillin-Clavulanic acid | 7 (8.4%) | 7 (8.4%) |
| Amikacin | 1 (1.2%) | 1 (1.2%) |
| Ceftazidime | 15 (18.1%) | 15 (18.1%) |
| Ciprofloxacin | 7 (8.4%) | 12 (14.5%) |
| Ceftriaxone | 15 (18.1%) | 15 (18.1%) |
| Ertapenem | 4 (4.8%) | 2 (2.4%) |
| Cefepime | 12 (14.5%) | 12 (14.5%) |
| Imipenem | 1 (1.2%) | 1 (1.2%) |
| Meropenem | 1 (1.2%) | 2 (2.4%) |
| Ofloxacin | 7 (8.4%) | 12 (14.5%) |
| Trimethoprim / sulfamethoxazole | 16 (19.3%) | 21 (25.3%) |
| Ticarcillin + clavulanic acid | 14 (16.9%) | 14 (16.9%) |
| Piperacillin + tazobactam | 6 (7.2%) | 5 (6.0%) |

<sup>a</sup>Vir-Kp: virulent *K. pneumoniae*
<sup>b</sup>Non-Vir-Kp: non-virulent *K. pneumoniae*

### Population structure and genomic insights into *K. pneumoniae* isolates

A total of 17,403 genes were identified in the collection, with 3,928 core genes (22.6%) conserved in at least 95% of the isolates. The remaining 77.4% accessory genes comprised 2,064 shell genes (11.9%) and 11,411 cloud genes (65.6%). Most accessory genes were rare and reported in <15% of isolates (S3 Fig). The pangenome analysis revealed a cluster of 12 *K. pneumoniae* possessing unique accessory gene acquisition (S1 Table). These organisms belonged to ST23 for sequence type and the K1 for capsular antigen type. For the somatic antigen, ten were O1, and the remaining two were O2afg. The isolates of this cluster were considered virulent, where nine had a virulence score of 5, one had 4, and the remaining two scored 3. The two highly virulent isolates (ERR2586355, ERR2586381) were MDR *K. pneumoniae* exhibiting resistance to β-lactams, fluoroquinolones, and sulfamethoxazole-trimethoprim.

The WGS analysis revealed at least 45 different STs with ST23 (16.7%) being the most prominent, followed by ST15 (8.3%), ST17 (5.6%), and ST25 (4.2%) (Fig 1). For LPS and O-antigen types, 36 isolates (50%) were O1, followed by serotypes O2afg, O3b, O2a and O4 present in 12.5%, 8.3%, 7.0% and 4.2% isolates, respectively. Overall, isolates of 16 different capsular serotypes were identified, predominantly K1 (15.3%), followed by K2 (8.3%), K10 (5.6%), K57 (5.6%) and K62 (5.6%). Based on ST, O and K serotyping, at least six different clusters of isolates were identified in the study population (clusters A, B, C, D, E and F). The largest cluster (A) had 12 isolates with a sequence type of ST23, and 83% of the members were O1 and K1 serotypes.

**Figure 1:**
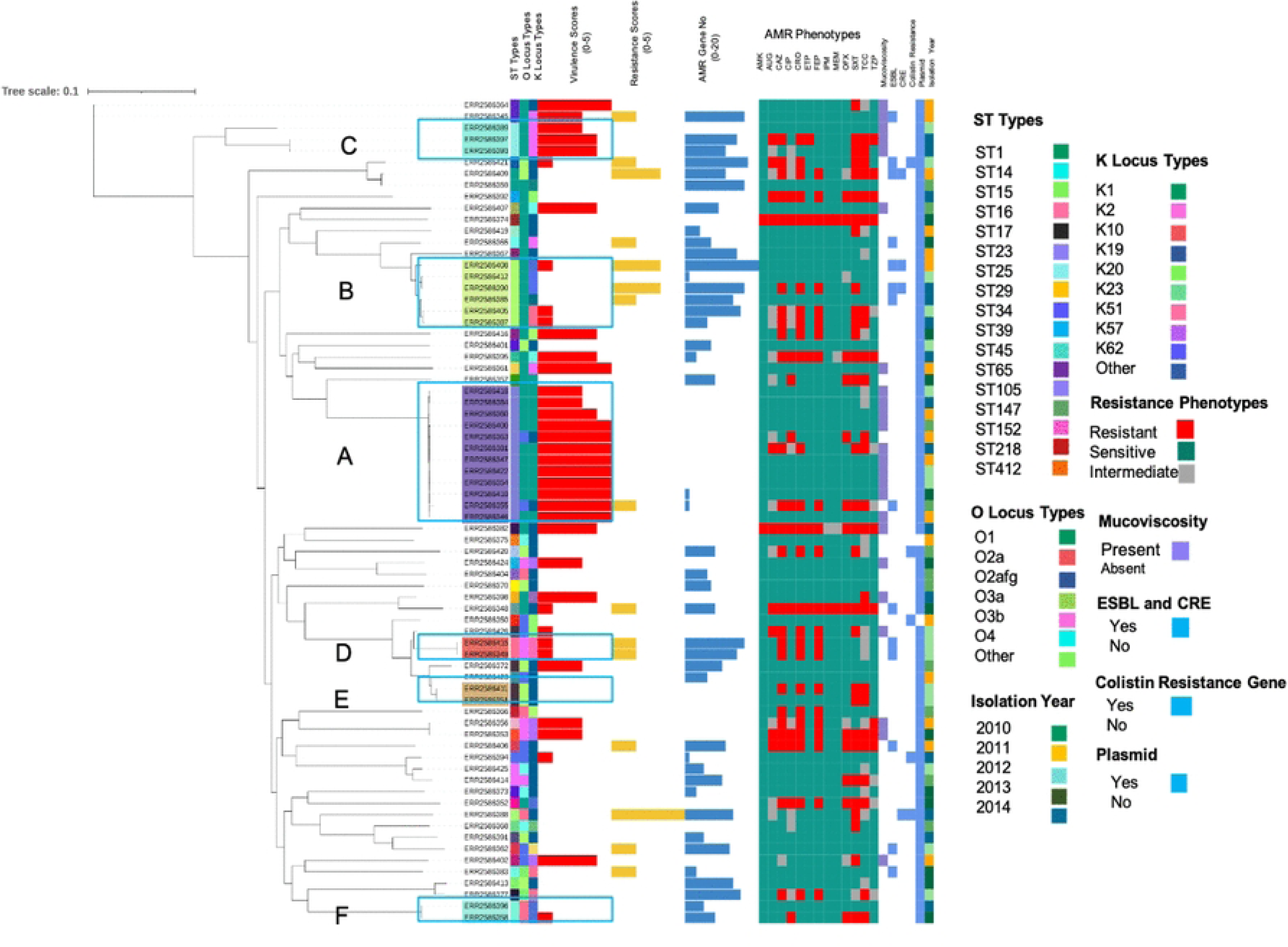
Core-genome phylogeny of *K. pneumoniae* study isolates. The SNPs-based phylogenetic tree as determined by Roary was constructed using the maximum likelihood method. The tree was produced by IQTREE and then visualised using iToL. Typing such as MLST, O and K typing, as well as virulence, resistance scoring and resistance gene calculation were performed in reference to Kleborate and Kaptive. AMR phenotypes were determined by the disc diffusion technique. The availability of genetic marker(s) was used to predict features like mucoviscosity, ESBL and CRE producer, presence of resistance and virulence plasmids and colistin resistance. The clustering of *K. pneumoniae* isolates was labelled accordingly (A, B, C, D, E).

For the virulence measurement of 72 isolates, 15.3%, 12.5%, and 11.1% isolates scored a virulence of 5, 4 and 3, respectively, whereas approximately half of the isolates (47.2%) scored 0. For AMR, approximately 80% of the isolates scored 0, meaning an absence of either ESBL or CRE or colistin resistance marker. At least ten different AMR genes (ARGs) conferring resistance to 7 to 11 diverse classes of antimicrobials were recorded in 27.8% of the isolates. According to genomic analysis, approximately 7% of the isolates were resistant to colistin, with three isolates showing mutation of the *pmrB* gene and two isolates acquiring the plasmid-mediated mobile *mcr-8.1* (Table 2). Approximately 40% of isolates had no ARGs. With the exception of two isolates, all other isolates (97.2%) harboured at least one plasmid (either virulence or resistance plasmids or both). The dominant plasmid types identified in the collection were repB_KLEB (18.5%), IncFIB(K) (16.7%), IncFIB(pKPHS1) (13.9%), and IncFII(K) (11.1%).

**Table 2:**
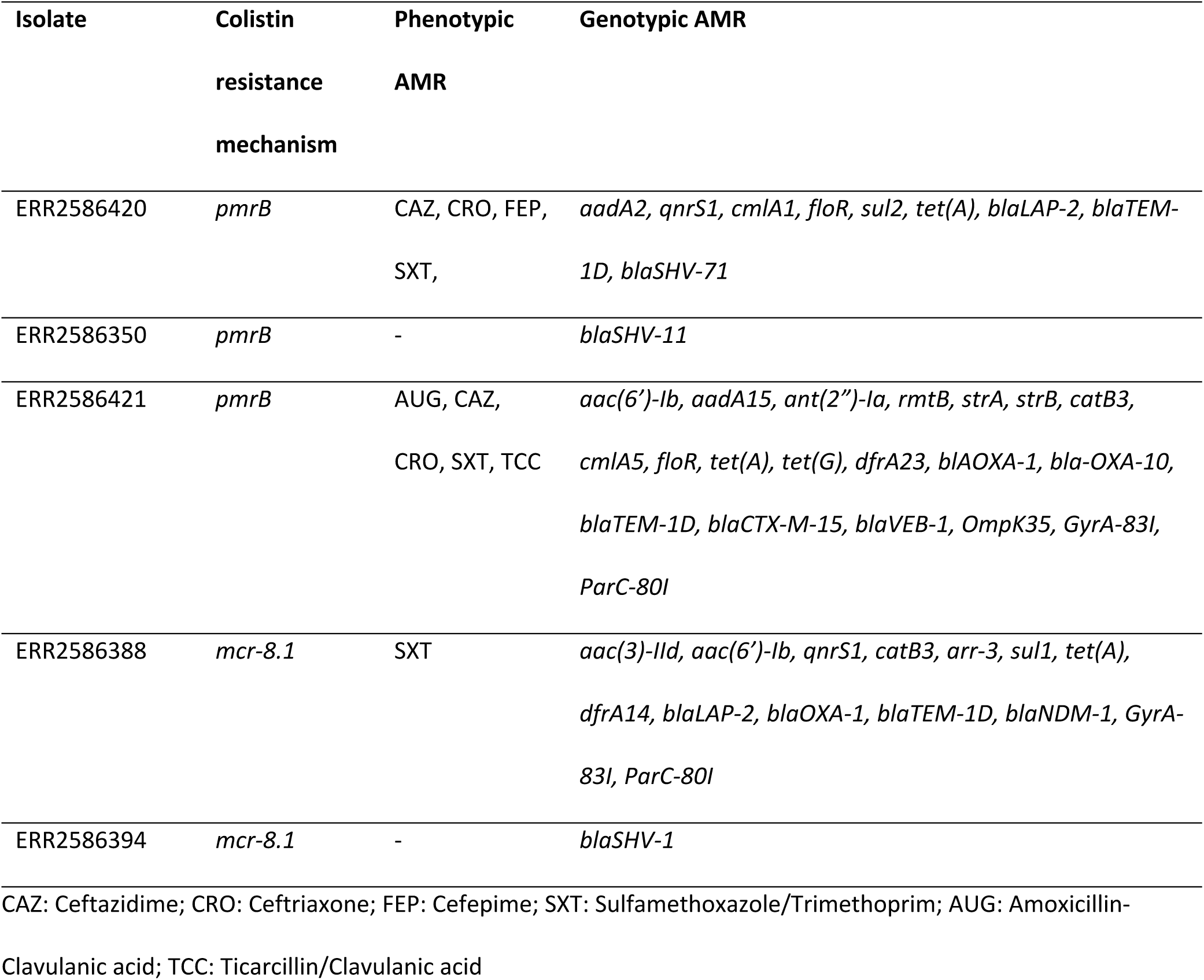
List of colistin resistance gene-positive *K. pneumoniae* isolates with phenotypic and genotypic AMR profile.

### Phenotypically colistin-resistant *K. pneumoniae* and LPS mass spectra

Out of eight phenotypically colistin-resistant isolates, one had an MIC of 64 mg/L, while the remaining ranged from 8 to 32 mg/L (Table 3). The native lipid A produced peaks at both m/z 1,824 and m/z 2,062 positions (±5 mass spectra unit) (S4 Fig). The ions at both m/z 1,824 and m/z 2,062 corresponded with bis-phosphorylated hexa- or hepta-acylated lipid A molecule with or without a hydroxyl group on the C’-2 fatty acyl chain. Concerning all eight phenotypically colistin-resistant *K. pneumoniae*, a further set of peaks was observed with a shift of m/z +131 mass unit at m/z 1,955 and m/z 2,193. These additional peaks indicate the presence of L-Ara4N on lipid A, induced by a chromosomal mutation (*pmrB*). The percentage of lipid A modification was >5% for all the tested isolates, indicating that they are all colistin-resistant. Among the isolates, the highest, 41.3% and the lowest, 8.3%, of L-Ara4N modification of lipid A were recorded. An increasing trend of a high percentage of lipid A modification was observed with the increase of the *K. pneumoniae* MIC value (correlation coefficient, r= 0.6 and coefficient of determination, R^2^= 0.4) (S5 Fig). *K. pneumoniae* that were phenotypically resistant to most antimicrobial classes recorded a high colistin MIC (32 mg/L to 64 mg/L) and possessed 2 to 22 genes conferring resistance to five to nine different categories of antimicrobials. Notably, no known colistin resistance markers were observed in any of the phenotypically colistin-resistant isolates.

**Table 3:** Results of MIC and MALDI-TOF tests on the phenotypically colistin-resistant *K. pneumoniae* isolates.

| Isolate | MIC (mg/L) | LPS Modification (%) | L-Ar4N Percentage | pENT Percentage | Phenotypic AMR profile by disc diffusion test | Genotypic AMR profile |
| --- | --- | --- | --- | --- | --- | --- |
| ERR2586370 | 16 | 15.7±0.8 | 100±0.0 | 0.0±0.0 | - | *strA, strB, catA1, sul2, tet(D), dfrA5, blaSHV-11* |
| ERR2586394 | 16 | 8.3±0.8 | 100±0.0 | 0.0±0.0 | - | *strA, strB, qnrS1, floR, sul1, sul2, tet(A), dfrA1, blaDHA-1, blaLAP-2, blaSHV-11* |
| ERR2586419 | 32 | 29.4±2.2 | 100±0.0 | 0.0±0.0 | SXT | *qnrS1, sul1, tet(A), dfrA1, blaDHA-1, blaSHV-186* |
| ERR2586404 | 8 | 12.0±4.2 | 100±0.0 | 0.0±0.0 | - | *aac(3)-IIa, aac(6')-Ib, aadA5, strA, strB, qnrB1, qnrB4, catB4, catA1, sul1, sul2, tet(A), dfrA1, dfrA14, blaDHA-1, blaTEM-1D, blaCTX-M-15, blaCTX-M-9, blaSHV-28, GyrA-83I, ParC-80I, blaOXA-1* |
| ERR2586344 | 16 | 28.6±2.9 | 100±0.0 | 0.0±0.0 | AUG, CAZ, CIP, CRO, OFX | *aac(6')-Ib, aadA15, aph3-Ia, qnrB4, qnrS1, mphA, catB3, arr-2, sul1, sul2, tet(A), dfrA14, blaDHA-1, blaLAP-2, blaOXA-1, blaTEM-1, blaSHV-1, GyrA-83I, ParC-80I* |
| ERR2586409 | 16 | 40.1±3.0 | 100±0.0 | 0.0±0.0 | CAZ, CRO, ETP, SXT, TCC, TZP | *aac(3)-IIa, aac(6'), strA, strB, catB4, sul2, dfrA14, blaOXA-1, blaTEM-1, blaCTX-M-15, blaSHV-28, GyrA-83I, ParC-80I* |
| ERR2586347 | 8 | 23.1±0.5 | 100±0.0 | 0.0±0.0 | - | *qnrS1, blaLAP-2, blaCTX-M-27* |
| ERR2586353 | 64 | 41.3±1.8 | 100±0.0 | 0.0±0.0 | AUG, CAZ, CIP, CRO, FEP, OFX, SXT, TCC, TZP | *aac(3)-IId, aac(6')-Ib, aadA16, strA, strB, qnrB4, qnrB6, mphA, catII, arr-3, sul1, sul2, tet(A), dfrA27, blaDHA-1, blaSHV-28, GyrA-83F, GyrA-87A, ParC-80I, aph3-Ia* |

SXT: Sulfamethoxazole/Trimethoprim; AUG: Amoxicillin-Clavulanic acid, CAZ: Ceftazidime; CIP: Ciprofloxacin; CRO: Ceftriaxone; OFX: Ofloxacin; ETP: Ertapenem, TCC:
Ticarcillin/Clavulanic acid, TZP: Piperacillin-Tazobactam; FEP: Cefepime

### Morphological changes of THP-1 due to LPS exposure

Initially, undifferentiated THP-1 cells had a regular, large, round, and single-cell morphology (S6 Fig). The density of cells shifted from 6.63 x 10^5^ per ml in the initial passages to 1.75 x 10^7^ per ml. Differentiated THP-1 cells were circular with dense nuclear material. With no evident clustering, the differentiated THP-1 cells were evenly distributed and adherent to the culture plate surface. With exposure to LPS, differentiated THP-1 cells showed observable death and displacement, giving a decreased cell density (S6 Fig).

### Type and degree of LPS modification affecting the inflammatory response

Between the two types of LPS modifications extracted from colistin-resistant *K. pneumoniae*, we observed that pEtN-conjugated LPS produced a significantly higher amount of IL-6, CXCL-8, and TNF-α compared to the L-Ara4N modification of LPS (*p*<0.05) (Fig 2).

**Fig 2:**
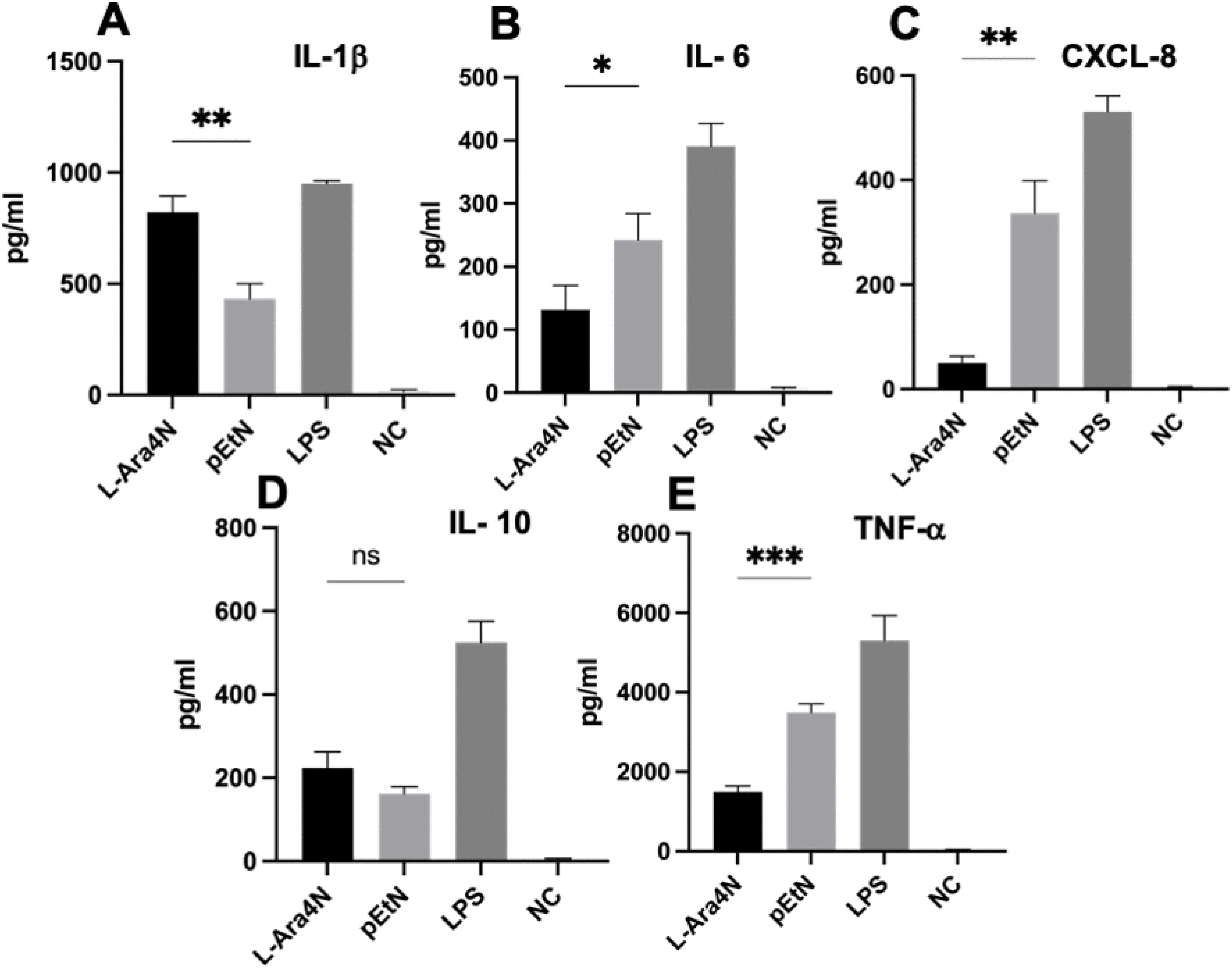
Immune response of differentiated THP-1 cells stimulated by modified LPS. Levels of IL-1β (A), IL-6 (B), CXCL-8 (C), IL-10 (D) and TNF-α (F) in cell culture supernatant after 24 hours of incubation by adding 10 ng/ml LPS having either L-Ara4N or pEtN modifications isolated from colistin-resistant *K. pneumoniae* isolates. Commercially available LPS (from *E. coli* O111:B4) and endotoxin-free water were used as positive and negative controls, respectively. Error bars represent SD (n=3). Student’s t-test was conducted to compare the LPS of L-Ara4N and pEtN. A *p*-value of ≤0.05 was accepted as statistically significant.

Conversely, interleukin-1 (IL-1β) production was significantly higher for L-Ara4N-LPS than pEtN-LPS (*p*=0.003) and no difference was observed in the production of IL-10 (*p*=0.067). Commercial unmodified LPS control triggered a consistently higher production of all these cytokines than the modified LPS extracted from the colistin-resistant study isolates.

To compare the impact of different degrees of LPS modification on immune response, two isolates were selected with 41.3% and 8.3% of lipid A modifications, denoted as “high” and “low” lipid A modification isolates, respectively. Inflammatory cytokine response from differentiated THP-1 cells revealed that the LPS with high lipid A modification triggered a greater immune response compared to low lipid A modification, as indicated by the greater levels of IL-1β, IL-6, and CXCL-8 (IL-8) (*p*<0.001) (Fig 3). On the contrary, IL-10 production was higher for the LPS with lower lipid A modification, although the difference was small (*p*=0.025). TNF-α production showed no substantial difference between low and high degrees of lipid A changes rendered by colistin resistance (*p*=0.177).

**Fig 3:**
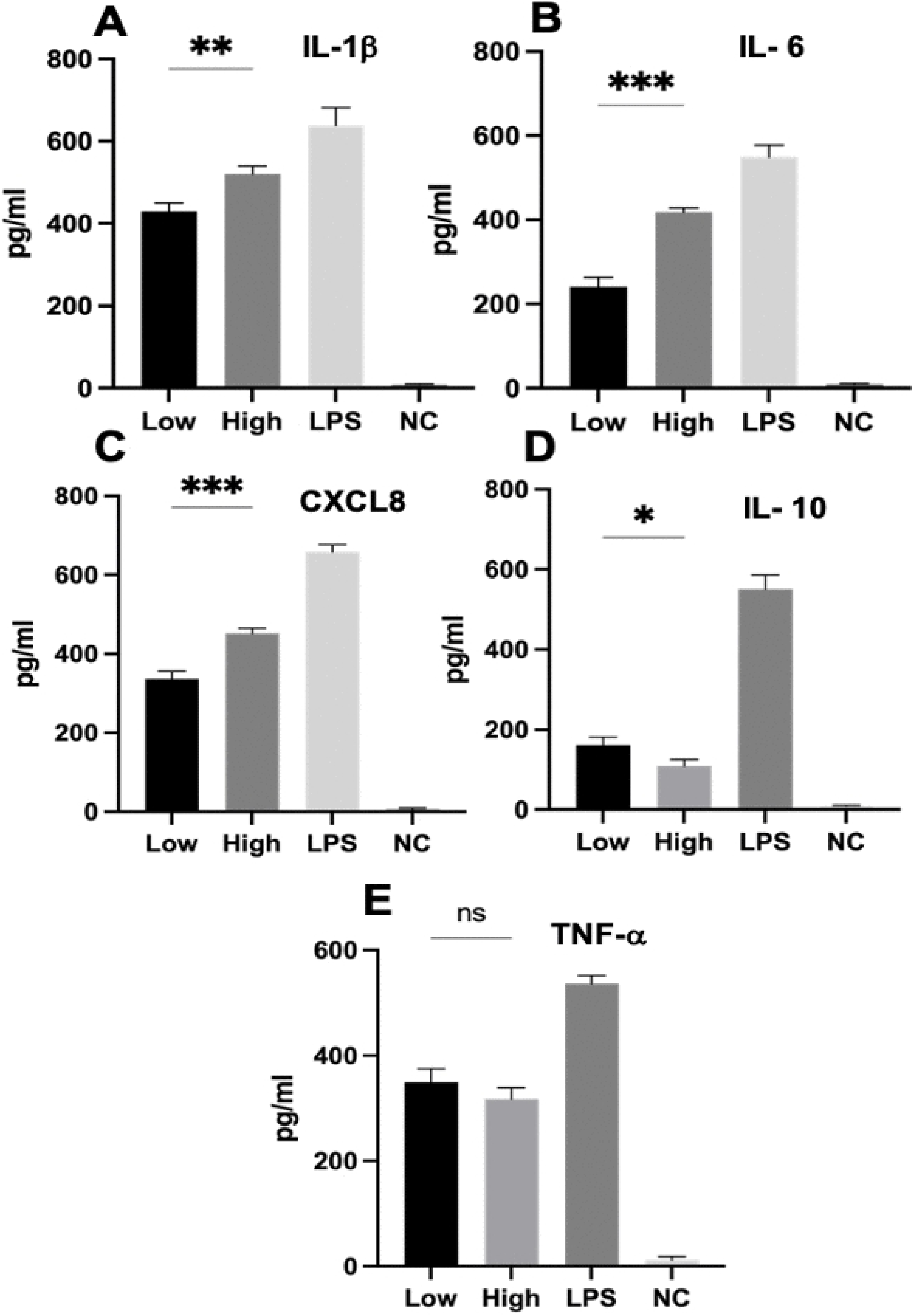
Expression of inflammatory cytokines by differentiated THP-1 cells. Levels of IL-1β (A), IL-6 (B), CXCL-8 (C), IL-10 (D) and TNF-α (F) in THP-cell supernatant after 24 hours of stimulation with 10 ng/ml LPS having two different percentages of lipid A modifications caused by colistin resistance. Commercially available LPS (from *E. coli* O111:B4) and endotoxin-free water were used as positive and negative controls, respectively. Data represent mean ± SD of triplicate. Student’s t-test was used to compare the high (41.3%) and low (8.3%) levels of LPS modifications. A *p-*value of ≤0.05 was accepted as statistically significant.

### LPS modification affecting colistin anti-endotoxin action

We selected two *K. pneumoniae* isolates with 28.6% and 15.7% LPS modifications, respectively. Colistin-treated LPS endotoxin was used to stimulate the cytokine response of differentiated THP-1 cells. Here, the isolate with 28.6% LPS modification triggered a significantly higher production of IL-1β, IL-6, IL-10, and TNF-α; the differential expression was notably high for IL-10 and TNF-α (*p*<0.001) (Fig 4). However, there was a higher level of CXCL-8 production for the isolate with 15.7% LPS modification. The commercial unmodified LPS control treated with colistin produced a significantly lower immune response than both experimental isolates.

**Fig 4:**
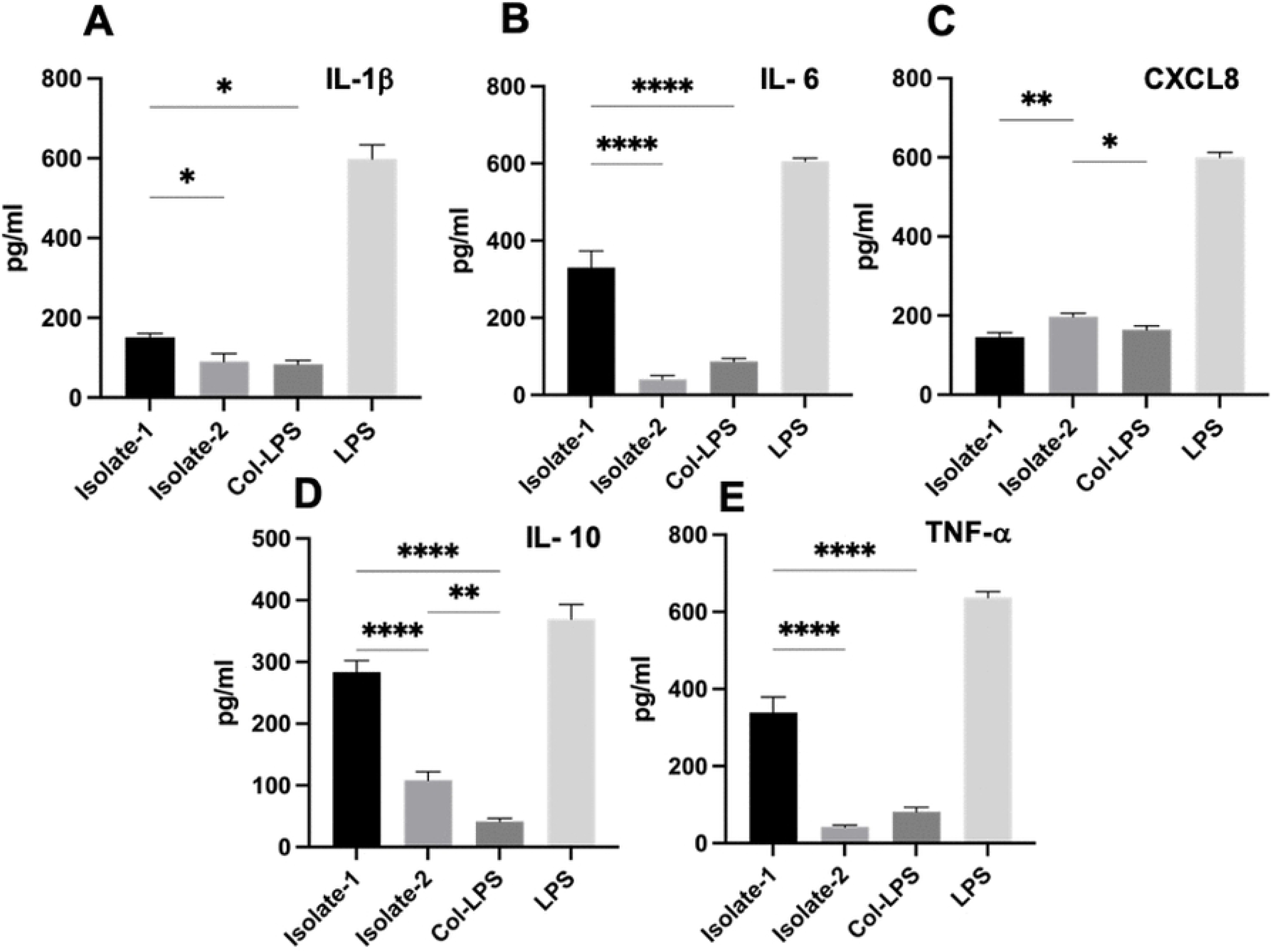
Result of the LPS neutralization test by colistin. Two colistin-resistant *K. pneumoniae* LPS treated with colistin were used to stimulate differentiated THP-1 cells. Here, A, B, C, D, and E plots represented IL-1β, IL-6, CXCL-8, IL-10 and TNF-α productions, respectively, for 3 replicates each. Commercially available LPS (from *E. coli* O111:B4) was used as a control. One-way ANOVA analysis indicated significant differences among isolates. A p-value of ≤0.05 marked statistical significance.

## Discussion

A convergence of hypervirulence and multidrug resistance makes *K. pneumoniae* a serious emerging cause of hospital-acquired infection. If such isolates acquire resistance to last-resort antimicrobials like colistin, then infections will result in grave consequences. Here, we identified eight phenotypically and five genotypically colistin-resistant clinical *K. pneumoniae* isolates in a tertiary healthcare centre in Vietnam. Two isolates were positive for plasmid-borne colistin resistance genes, *mcr-8.1*. Since *mcr* genes are highly horizontally transmissible, seeing an unprecedented rise in the rate of colistin resistance in the last decade among nosocomial pathogens is not surprising [28]. Among the five genotypically colistin-resistant isolates identified in this study, three had a mutation in the *mgrB* gene, the negative regulator of the PhoPQ two-component systems [29,30]. Though resistance to colistin can be acquired either through chromosomal mutation or plasmid-borne mechanisms, in *K. pneumoniae*, chromosome-encoded resistance is more prominent than *mcr* genes [31]. Large-scale screening of clinical colistin-resistant *K. pneumoniae* isolates showed that in *K. pneumoniae,* colistin resistance is mainly mediated by the alteration or disruption of the *mgrB* gene [11,29]. Studies have established that interruption of the *mgrB* gene by different insertion sequences, such as *IS10-like* and *IS5-like* elements, is common in *K. pneumoniae* [32,33]. Alternatively, one of the plasmid-mediated colistin resistance genes, *mcr-8.1*, identified in two isolates of this study, was first reported in *K. pneumoniae* [34]. Besides *mcr-8.1*, other *mcr* genes (*mcr-1*, *mcr-3* and *mcr-7*) have also been reported in *K. pneumoniae* but were not identified here [7,35,36].

Since colistin is the last option to treat multidrug-resistant infections, the rapid detection of colistin resistance is of the utmost importance in clinical settings. In this study, the recently validated MALDIxin test was applied to detect colistin resistance within 30 minutes using mass spectrum data [11]. The motivation behind applying the MALDIxin test, in addition to the phenotypic resistance profile of the organism, was that it also provides data regarding the genetic basis of the colistin resistance phenotype. Mass spectrum data enables the differentiation of chromosomal mutations from the plasmid-borne mechanisms. This test has been successfully optimised on a routine MALDI Biotyper Sirius system for quick colistin resistance detection for important members of the Enterobacteriaceae family, such as *E. coli*, *Salmonella* spp., *Acinetobacter baumannii*, and *K. pneumoniae* [37–39].

The degree of lipid A modification, as revealed by the MALDIxin test in colistin-resistant isolates of this study, generated a positive correlation with MIC values. A trend of a higher percentage of modified lipid A was observed with a high *K. pneumoniae* MIC value against colistin. At least a 5% modification of the total LPS on the outer membrane of the Gram-negative bacteria is required to show phenotypic colistin resistance in *K. pneumoniae* [11]. Further studies can be carried out to check the correlation between different MIC values against colistin and the degree of LPS modification in the MDR organism. Nevertheless, seven isolates showing colistin resistance based on MALDIxin data did not account for any colistin resistance gene reported to date. Reduced susceptibility or resistance to colistin without the presence of any known mechanism has been reported [14]. These findings necessitate the exploration of the colistin-resistant *K. pneumoniae* isolates to understand the genetic basis of their resistance. Colistin resistance gene-positive *K. pneumoniae* showing a phenotypically susceptible profile indicates that a single gene may not be sufficient to express the genetic trait of colistin resistance [40].

Since lipid A is a potent immune stimulator, we hypothesised that the type and degree of LPS modification triggered by colistin resistance in *K. pneumoniae* may affect its immunogenic property. We observed that, except for IL-1β, a pEtN modification of LPS elicited a greater immune response to IL-6, CXCL-8 and TNF-α than the L-Ara4N LPS modification. There are limited or no data available directly comparing the immune response between L-Ara4N and pEtN conjugated LPS. *E. coli* isolates expressing the *mcr-3* gene, coding for pEtN, trigger macrophage cells for the elevated production of IL-6, IL-1β and TNF-α compared to the mutant isolate [13]. A potential reason behind this observation may be the enhanced recognition of the pEtN group of modified LPS by the host TLR4 [41]. Nevertheless, contradictory findings also exist where *mcr-1* expression is reported to be associated with lower inflammatory response for IL-6, TNF-α [16]. A low IL-1β production was observed in *mcr-1*-positive *E. coli* coding pEtN compared to the *mcr-1*-negative parental strain [42]. In relation to the degree of LPS modification, we observed that the higher degree of LPS modification caused by colistin resistance induced a greater immune signal in the THP-1 cells in producing IL-1β, IL-6, and CXCL-8 markers. The number of phosphoryl substituents, such as pEtN or phosphate groups on lipid A, positively correlates with the activation of the TLR4-myeloid differentiation factor 2 (TLR4-MD2) pathway [43]. However, the reason(s) for a different outcome for IL-10 and TNF-α are not clear. While comparing different LPS modifications for inflammatory responses, the type of modification (L-Ara4N or pEtN) should also be considered to get a better holistic understanding.

We further evaluated the effect of LPS modification on colistin anti-endotoxin function. A high degree of LPS modification disrupts endotoxin neutralisation by colistin, impacting the clinical management of septic shock cases [44,45]. In sepsis and septic shock patients, polymyxin B immobilised fibre column (PMX) is commonly used for endotoxin removal and reduction of blood inflammatory cytokine levels such as IL-6, IL-17A, and TNF-α [46]. Our findings indicate that the LPS modification of *K. pneumoniae* caused by colistin resistance may affect the outcome of this therapy. Further studies should be conducted to measure the impact of different LPS modifications on the effectiveness of PMX therapy, especially using isolates with different degrees and types (L-Ara4N or pEtN) of LPS modifications.

## Conclusions

We elucidated the impact of colistin resistance-mediated LPS modification of *K. pneumoniae* on the inflammatory response in THP-1-derived macrophages. Our findings suggest that the immune trigger depends on the type and degree of the LPS modifications. Furthermore, this modification can also potentially affect the colistin anti-endotoxin function. These findings provide evidence of altered host-microbe interaction dynamics in the context of colistin resistance.

## Acknowledgements

We thank the team members of the Hospital for Tropical Diseases (HTD), Vietnam, for providing the *K. pneumoniae* isolates.

## Supporting information

**S1 Table:** List of unique genes identified in the pangenome cluster of 12 *K. pneumoniae* isolates, along with their respective function

**S1 Fig:** Sketch of the optimised MALDIxin test to detect colistin resistance in *K. pneumoniae*

**S2 Fig:** Outline of the workflow of THP-1 cell culture to measure the impact of colistin resistance-mediated LPS modification on host immunity

**S3 Fig:** Soft-core and accessory genome composition of the *K. pneumoniae* isolates

The pan genome matrix, showing the presence/absence of each gene in the strain. The matrix columns represent all the genes in the pan genome, and the blue colour represents the strains in which they are present.

**S4 Fig:** Representative mass spectra of naive and modified lipid A of *K. pneumoniae*

**S5 Fig:** Correlation between minimum inhibitory concentration (MIC) value and percentage of colistin-mediated lipid A modification

**S6 Fig:** Effect of LPS on differentiated THP-1 cells

(A) Undifferentiated THP-1 cells, (B) Differentiated PMA/THP-1 cells and (C) PMA/THP-1 cells after 24 hours of LPS exposure. Images were captured using a light microscope at 100x magnification.

## References

1. Chong PM, McCorrister SJ, Unger MS, Boyd DA, Mulvey MR, Westmacott GR. MALDI-TOF MS detection of carbapenemase activity in clinical isolates of *Enterobacteriaceae* spp., *Pseudomonas aeruginosa*, and *Acinetobacter baumannii* compared against the Carba-NP assay. J Microbiol Methods. 2015;111: 21–23. doi:10.1016/j.mimet.2015.01.024

2. Gatin L, Mghir AS, Mouton W, Laurent F, Ghout I, Rioux-Leclercq N, et al. Colistin-containing cement spacer for treatment of experimental carbapenemase-producing *Klebsiella pneumoniae* prosthetic joint infection. Int J Antimicrob Agents. 2019;54: 456–462. doi:10.1016/j.ijantimicag.2019.07.009

3. Roberts JL, Cattoz B, Schweins R, Beck K, Thomas DW, Griffiths PC, et al. In Vitro Evaluation of the Interaction of Dextrin–Colistin Conjugates with Bacterial Lipopolysaccharide. J Med Chem. 2016;59: 647–654. doi:10.1021/acs.jmedchem.5b01521

4. Narimisa N, Goodarzi F, Bavari S. Prevalence of colistin resistance of *Klebsiella pneumoniae* isolates in Iran: a systematic review and meta-analysis. Ann Clin Microbiol Antimicrob. 2022;21: 29. doi:10.1186/s12941-022-00520-8

5. Liu Y, Lin Y, Wang Z, Hu N, Liu Q, Zhou W, et al. Molecular Mechanisms of Colistin Resistance in *Klebsiella pneumoniae* in a Tertiary Care Teaching Hospital. Front Cell Infect Microbiol. 2021;11: 673503. doi:10.3389/fcimb.2021.673503

6. Leung LM, Cooper VS, Rasko DA, Guo Q, Pacey MP, McElheny CL, et al. Structural modification of LPS in colistin-resistant, KPC-producing *Klebsiella pneumoniae*. J Antimicrob Chemother. 2017;72: 3035–3042. doi:10.1093/jac/dkx234

7. Chen F-J, Lauderdale T-L, Huang W-C, Shiau Y-R, Wang H-Y, Kuo S-C. Emergence of *mcr-1*, *mcr-3* and *mcr-8* in clinical *Klebsiella pneumoniae* isolates in Taiwan. Clin Microbiol Infect. 2021;27: 305–307. doi:10.1016/j.cmi.2020.07.043

8. Karki D, Dhungel B, Bhandari S, Kunwar A, Joshi PR, Shrestha B, et al. Antibiotic resistance and detection of plasmid mediated colistin resistance *mcr-1* gene among *Escherichia coli* and *Klebsiella pneumoniae* isolated from clinical samples. Gut Pathog. 2021;13: 45. doi:10.1186/s13099-021-00441-5

9. Llobet E, Martínez-Moliner V, Moranta D, Dahlström KM, Regueiro V, Tomás A, et al. Deciphering tissue-induced *Klebsiella pneumoniae* lipid A structure. Proc Natl Acad Sci U S A. 2015;112: E6369–6378. doi:10.1073/pnas.1508820112

10. Clements A, Tull D, Jenney AW, Farn JL, Kim S-H, Bishop RE, et al. Secondary acylation of *Klebsiella pneumoniae* lipopolysaccharide contributes to sensitivity to antibacterial peptides. J Biol Chem. 2007;282: 15569–15577. doi:10.1074/jbc.M701454200

11. Dortet L, Broda A, Bernabeu S, Glupczynski Y, Bogaerts P, Bonnin R, et al. Optimization of the MALDIxin test for the rapid identification of colistin resistance in *Klebsiella pneumonia*e using MALDI-TOF MS. J Antimicrob Chemother. 2020;75: 110–116. doi:10.1093/jac/dkz405

12. Gauthier AE, Rotjan RD, Kagan JC. Lipopolysaccharide detection by the innate immune system may be an uncommon defence strategy used in nature. Open Biol. 12: 220146. doi:10.1098/rsob.220146

13. Yin W, Ling Z, Dong Y, Qiao L, Shen Y, Liu Z, et al. Mobile colistin resistance enzyme MCR-3 facilitates bacterial evasion of host phagocytosis. Adv Sci. 2021;8: 2101336. doi:10.1002/advs.202101336

14. Avendaño-Ortiz J, Ponce-Alonso M, Llanos-González E, Barragán-Prada H, Barbero-Herranz R, Lozano-Rodríguez R, et al. The impact of colistin resistance on the activation of innate immunity by lipopolysaccharide modification. Infect Immun. 2023;91: e00012–23. doi:10.1128/iai.00012-23

15. Hollaus R, Ittig S, Hofinger A, Haegman M, Beyaert R, Kosma P, et al. Chemical synthesis of Burkholderia Lipid A modified with glycosyl phosphodiester-linked 4-amino-4-deoxy-β-L-arabinose and its immunomodulatory potential. Chem Weinh Bergstr Ger. 2015;21: 4102–4114. doi:10.1002/chem.201406058

16. Yang Q, Li M, Spiller OB, Andrey DO, Hinchliffe P, Li H, et al. Balancing *mcr-1* expression and bacterial survival is a delicate equilibrium between essential cellular defence mechanisms. Nat Commun. 2017;8: 2054. doi:10.1038/s41467-017-02149-0

17. Kidd TJ, Mills G, Sá-Pessoa J, Dumigan A, Frank CG, Insua JL, et al. A *Klebsiella pneumoniae* antibiotic resistance mechanism that subdues host defences and promotes virulence. EMBO Mol Med. 2017;9: 430–447. doi:10.15252/emmm.201607336

18. Rodrigues C, Passet V, Rakotondrasoa A, Brisse S. Identification of *Klebsiella pneumoniae*, *Klebsiella quasipneumoniae*, *Klebsiella variicola* and related phylogroups by MALDI-TOF mass spectrometry. Front Microbiol. 2018;9. doi:10.3389/fmicb.2018.03000

19. CLSI. Performance standards for antimicrobial susceptibility testing. 36^th^ ed. Clinical and Laboratory Standards Institute supplement M100. CLSI, Wayne, PA, USA. 2026.

20. Roberts JL, Cattoz B, Schweins R, Beck K, Thomas DW, Griffiths PC, et al. In vitro evaluation of the interaction of Dextrin–Colistin conjugates with bacterial lipopolysaccharide. J Med Chem. 2016;59: 647–654. doi:10.1021/acs.jmedchem.5b01521

21. Lam MMC, Wick RR, Watts SC, Cerdeira LT, Wyres KL, Holt KE. A genomic surveillance framework and genotyping tool for *Klebsiella pneumoniae* and its related species complex. Nat Commun. 2021;12: 4188. doi:10.1038/s41467-021-24448-3

22. Seemann T. Prokka: rapid prokaryotic genome annotation. Bioinformatics. 2014;30: 2068–2069. doi:10.1093/bioinformatics/btu153

23. Page AJ, Cummins CA, Hunt M, Wong VK, Reuter S, Holden MTG, et al. Roary: rapid large-scale prokaryote pan genome analysis. Bioinformatics. 2015;31: 3691–3693. doi:10.1093/bioinformatics/btv421

24. Di Tommaso P, Chatzou M, Floden EW, Barja PP, Palumbo E, Notredame C. Nextflow enables reproducible computational workflows. Nat Biotechnol. 2017;35: 316–319. doi:10.1038/nbt.3820

25. Underwood A. GHRU (Genomic Surveillance of Antimicrobial Resistance) Retrospective 1 Bioinformatics Methods V.4. 2020. doi.10.17504/protocols.io.bp2l6b11kgqe/v4

26. Minh BQ, Schmidt HA, Chernomor O, Schrempf D, Woodhams MD, von Haeseler A, et al. IQ-TREE 2: New models and efficient methods for phylogenetic inference in the genomic era. Mol Biol Evol. 2020;37: 1530–1534. doi:10.1093/molbev/msaa015

27. Letunic I, Bork P. Interactive tree of life (iTOL) v3: an online tool for the display and annotation of phylogenetic and other trees. Nucleic Acids Res. 2016;44: W242–W245. doi:10.1093/nar/gkw290

28. Andrade FF, Silva D, Rodrigues A, Pina-Vaz C. Colistin update on its mechanism of action and resistance, present and future challenges. Microorganisms. 2020;8: 1716. doi:10.3390/microorganisms8111716

29. Aires CAM, Pereira PS, Asensi MD, Carvalho-Assef APD. mgrB mutations mediating Polymyxin B resistance in *Klebsiella pneumoniae* isolates from rectal surveillance swabs in Brazil. Antimicrob Agents Chemother. 2016;60: 6969–6972. doi:10.1128/AAC.01456-16

30. Olaitan AO, Morand S, Rolain J-M. Mechanisms of polymyxin resistance: acquired and intrinsic resistance in bacteria. Front Microbiol. 2014;5: 643. doi:10.3389/fmicb.2014.00643

31. Poirel L, Jayol A, Nordmann P. Polymyxins: Antibacterial activity, susceptibility testing, and resistance mechanisms encoded by plasmids or chromosomes. Clin Microbiol Rev. 2017;30: 557–596. doi:10.1128/CMR.00064-16

32. Cannatelli A, Giani T, D’Andrea MM, Di Pilato V, Arena F, Conte V, et al. MgrB inactivation is a common mechanism of colistin resistance in KPC-producing *Klebsiella pneumoniae* of clinical origin. Antimicrob Agents Chemother. 2014;58: 5696–5703. doi:10.1128/aac.03110-14

33. Poirel L, Jayol A, Bontron S, Villegas M-V, Ozdamar M, Türkoglu S, et al. The mgrB gene as a key target for acquired resistance to colistin in *Klebsiella pneumoniae*. J Antimicrob Chemother. 2015;70: 75–80. doi:10.1093/jac/dku323

34. Wang X, Wang Y, Zhou Y, Li J, Yin W, Wang S, et al. Emergence of a novel mobile colistin resistance gene, *mcr-8*, in NDM-producing *Klebsiella pneumoniae*. Emerg Microbes Infect. 2018;7: 1–9. doi:10.1038/s41426-018-0124-z

35. Yang Y-Q, Li Y-X, Lei C-W, Zhang A-Y, Wang H-N. Novel plasmid-mediated colistin resistance gene *mcr-7.1* in *Klebsiella pneumoniae*. J Antimicrob Chemother. 2018;73: 1791– 1795. doi:10.1093/jac/dky111

36. Di Pilato V, Arena F, Tascini C, Cannatelli A, Henrici De Angelis L, Fortunato S, et al. *mcr-1.2*, a new *mcr* variant carried on a transferable plasmid from a colistin-resistant KPC carbapenemase-producing *Klebsiella pneumoniae* strain of sequence type 512. Antimicrob Agents Chemother. 2016;60: 5612–5615. doi:10.1128/aac.01075-16

37. Dortet L, Bonnin RA, Pennisi I, Gauthier L, Jousset AB, Dabos L, et al. Rapid detection and discrimination of chromosome- and MCR-plasmid-mediated resistance to polymyxins by MALDI-TOF MS in *Escherichia coli*: the MALDIxin test. J Antimicrob Chemother. 2018;73: 3359–3367. doi:10.1093/jac/dky330

38. Dortet L, Bonnin RA, Le Hello S, Fabre L, Bonnet R, Kostrzewa M, et al. Detection of colistin resistance in *Salmonella enterica* using MALDIxin test on the routine MALDI Biotyper Sirius Mass Spectrometer. Front Microbiol. 2020;11: 1141. doi:10.3389/fmicb.2020.01141

39. Dortet L, Potron A, Bonnin RA, Plesiat P, Naas T, Filloux A, et al. Rapid detection of colistin resistance in *Acinetobacter baumannii* using MALDI-TOF-based lipidomics on intact bacteria. Sci Rep. 2018;8: 16910. doi:10.1038/s41598-018-35041-y

40. Macesic N, Blakeway LV, Stewart JD, Hawkey J, Wyres KL, Judd LM, et al. Silent spread of mobile colistin resistance gene *mcr-9.1* on IncHI2 ‘superplasmids’ in clinical carbapenem-resistant Enterobacterales. Clin Microbiol Infect. 2021;27: 1856.e7–1856.e13. doi:10.1016/j.cmi.2021.04.020

41. Cullen TW, O’Brien JP, Hendrixson DR, Giles DK, Hobb RI, Thompson SA, et al. EptC of *Campylobacter jejuni* mediates phenotypes involved in host interactions and virulence. Infect Immun. 2013;81: 430–440. doi:10.1128/iai.01046-12

42. Mattiuz G, Nicolò S, Antonelli A, Giani T, Baccani I, Cannatelli A, et al. *mcr-1* gene expression modulates the inflammatory response of human macrophages to *Escherichia coli*. Infect Immun. 2020;88: 10.1128/iai.00018-20. doi:10.1128/iai.00018-20

43. Liu M, John CM, Jarvis GA. Phosphoryl moieties of Lipid A from *Neisseria meningitidis* and *N. gonorrhoeae* lipooligosaccharides play an important role in activation of both MyD88- and TRIF-dependent TLR4–MD-2 signaling pathways. J Immunol. 2010;185: 6974–6984. doi:10.4049/jimmunol.1000953

44. Shoji H, Opal SM. Therapeutic rationale for endotoxin removal with Polymyxin B immobilized fiber column (PMX) for septic shock. Int J Mol Sci. 2021;22: 2228. doi:10.3390/ijms22042228

45. Hussein MH, Kato T, Sugiura T, Daoud GA, Suzuki S, Fukuda S, et al. Effect of hemoperfusion using Polymyxin B-immobilized fiber on IL-6, HMGB-1, and IFN gamma in a neonatal sepsis model. Pediatr Res. 2005;58: 309–314. doi:10.1203/01.PDR.0000169995.25333.F7

46. Mitaka C, Kusaoi M, Kawagoe I, Satoh D. Up-to-date information on polymyxin B-immobilized fiber column direct hemoperfusion for septic shock. Acute Crit Care. 2021;36: 85–91. doi:10.4266/acc.2021.00150

